# Effective strategies for preventing reestablishment of malaria in areas with recent elimination and high transmission potential

**DOI:** 10.1101/766048

**Authors:** Jaline Gerardin, Caitlin A. Bever, Daniel Bridenbecker, Thomas P. Eisele, Busiku Hamainza, John M. Miller, Edward A. Wenger

## Abstract

Maintaining zero transmission after malaria elimination will be a challenging task for many countries where malaria is still endemic. When local transmission potential is high, and importation of malaria infections continues from neighboring areas with ongoing transmission, malaria programs must develop robust surveillance and outbreak response systems. However, the requirements for such systems remain unclear. Using an agent-based, spatial microsimulation model of two areas in southern Zambia, where elimination efforts are currently underway, we compare the ability of various routine and reactive intervention packages to maintain near-zero prevalence in the face of continued importation. We find that in formerly moderate-transmission areas, high treatment rate of symptomatic malaria is sufficient to prevent reestablishment of malaria. Routine redistributions of insecticide-treated nets and reactive case detection with antimalarial drugs cannot completely compensate for inadequate case management. In formerly high-transmission areas, excellent case management and maintenance of good bednet coverage are both required to prevent resurgence, and outbreak response with antimalarial drugs or additional vector control is also necessary. These results begin to describe the essential criteria for operations that successfully prevent reestablishment of malaria post-elimination and highlight the need for both long-term, sustainable excellence in primary care and comprehensive surveillance that feeds into rapid and flexible outbreak response.

**Author Summary:** The global community is working toward malaria elimination, but some areas will eliminate before others. Eliminated areas will need to develop intervention programs capable of preventing imported infections from leading to reestablishment, a particular challenge when transmission was previously very high. Past experience has shown that stopping elimination interventions leads to massive resurgence, but it is unclear which interventions must be continued, which can be stopped to conserve resources, and what new interventions should be deployed. Using a simulation model built to capture malaria transmission and intervention history of two areas that recently made enormous progress toward elimination, we tested how well different intervention programs were able to prevent reestablishment of malaria. We found that treating as many symptomatic cases as possible was the single most important intervention to implement. In some contexts, this intervention alone was sufficient to prevent reestablishment. Other areas with historically higher transmission required maintaining vector control to contain mosquito populations. Localized outbreak response with antimalarial drugs or additional vector control was also necessary and predicted to be a highly efficient use of resources. These findings provide quantitative guidance for policy-makers considering how to stratify eliminated areas and plan new operational modes for the post-elimination era.

## Introduction

Malaria burden has declined tremendously over the last two decades, and more countries are now malaria-free than ever before (1, 2). Global policymakers have issued technical guidance for countries on how to accelerate their progress toward malaria elimination and how to maintain elimination once it has been reached (3-5). Guidance around maintaining elimination has focused on receptivity, the degree to which a local area allows for the transmission of malaria parasites from a human through a vector mosquito to another human, and importation risk, the risk of potential influx of parasites into an area via infected individuals or infected vectors. Together, these quantities describe the likelihood of local transmission post-elimination. However, current guidance has limited quantitative metrics around distinguishing areas with high and low receptivity or high and low importation rate, and recommendations for what operations to implement under different conditions of receptivity and importation rate are therefore difficult to apply.

Most of the countries that have already eliminated have lower receptivity, and they maintain elimination through excellent clinical surveillance, focal case investigation, and in some cases entomological surveillance and response (6). In contrast, many of the remaining malaria-endemic countries have much higher intrinsic transmission potential and are surrounded by other countries facing the same issues, making their path toward elimination technically, operationally, and financially challenging. Strategies to prevent reestablishment of malaria in these formerly high-transmission areas must be developed, especially if cases continue to be imported from neighbors that have not yet eliminated. Previous experience in the Garki area of northern Nigeria, where indoor residual spraying (IRS) of insecticides and mass drug administration (MDA) with antimalarials briefly brought a few villages to elimination or near-elimination in the 1970s, showed that transmission could rapidly bounce back to pre-intervention levels after ceasing intervention activities (7). Uganda has experienced similar resurgence of malaria following cessation of IRS (8).

These observations raise the question of how to maintain elimination, or near-elimination, in areas with formerly high transmission. Key operational questions, including the intensity of clinical surveillance necessary, how long to continue routine vector control, what kind of outbreak response to implement, and even whether to maintain the routine interventions that achieved elimination or to move toward a more reactive operational framework, remain unanswered.

Mathematical modeling and microsimulations have been used to design intervention packages for malaria control and elimination and to understand the roles interventions play alone and in combination to achieve programmatic goals (9-14). A theoretical exploration predicted that malaria elimination should be “sticky” in the sense that an eliminated area should be able to maintain elimination as long as treatment rates for symptomatic cases were high (15), while other work in a microsimulation model found that maintaining vector control would also be necessary in most areas, especially under higher importation rates (16). Previous work in a susceptible-infectious-susceptible (SIS) model also underlined the role of case management, but added that reactive case detection (RCD), where households surrounding index cases also receive intervention, could be helpful for maintaining elimination (17).

While previous studies on prevention of reestablishment have explored this problem in abstracted and non-spatial frameworks, spatial epidemiological models are well-positioned to describe how imported infections can spread and how reactive interventions may or may not succeed in containing transmission. In this study, we deploy two spatial microsimulation models of specific settings in southern Zambia, parametrized with rich study data from the area, to compare the roles of case management, routine vector control, reactive distribution of antimalarial drugs, and reactive vector control in maintaining near-zero prevalence in areas that have recently eliminated malaria.

## Results

To investigate what interventions are necessary to prevent re-establishment of malaria in settings that have recently interrupted transmission, we constructed models of two areas in Southern Province, Zambia, where intensive intervention programs have greatly reduced malaria burden over the last few years (18-20) (Figure 1A). We focus on two health facility catchment areas (HFCAs) with distinct malaria ecologies and intervention histories, Bbondo and Luumbo HFCAs. Keeping each HFCA’s history intact, including population immunity levels, we instantaneously remove all parasites from both humans and mosquitoes, then observe the HFCA over the next ten years as residents continue to travel to areas with ongoing transmission and return home with malaria infection (Figure 1B). During the tenth year, we assess the mean prevalence of any malaria infection in all human residents of the HFCA (Figure 1C). Successful intervention packages are those that maintain prevalence near zero or at very low levels.

**Figure 1.**
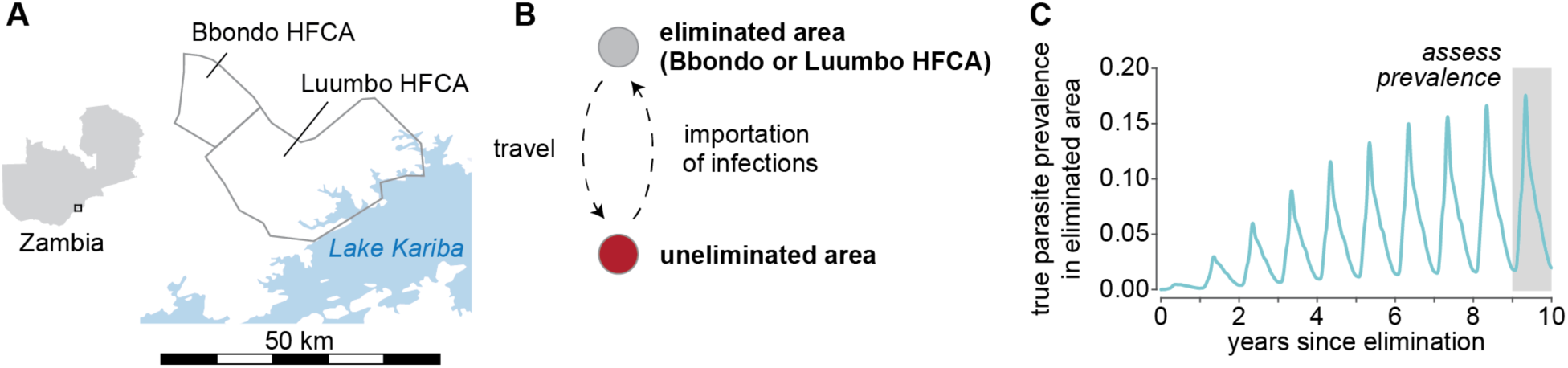
Health facility catchment areas in Zambia used as model systems for testing intervention programs for preventing reestablishment of malaria after local elimination. (A) Bbondo and Luumbo HFCAs are in the Lake Kariba area of Southern Province, Zambia. (B) Residents of simulated Bbondo or Luumbo, where malaria was artificially eliminated, travel to and from a nearby area where malaria transmission is ongoing, bringing back imported infections to their home HFCA. (C) Various intervention scenarios are simulated for 10 years, and prevalence of malaria infection is assessed in the 10^th^ year to compare the efficacy of various interventions in preventing resurgence. Trace shown is true prevalence in Bbondo under 12 importations per year, 50% case management, rfTAT at the household level with 100% response rate, mean of 100 stochastic realizations.

### Preventing resurgence in areas of formerly moderate transmission requires good case management

Bbondo HFCA contains around 745 households and 8,000 residents (19) (Figure 2A). Historically, transmission was moderate, with prevalence by rapid diagnostic test (RDT) of 9% in the dry season. In addition to the health clinic, six community health workers (CHWs) are active in the area, with one collocated with the clinic.

**Figure 2.**
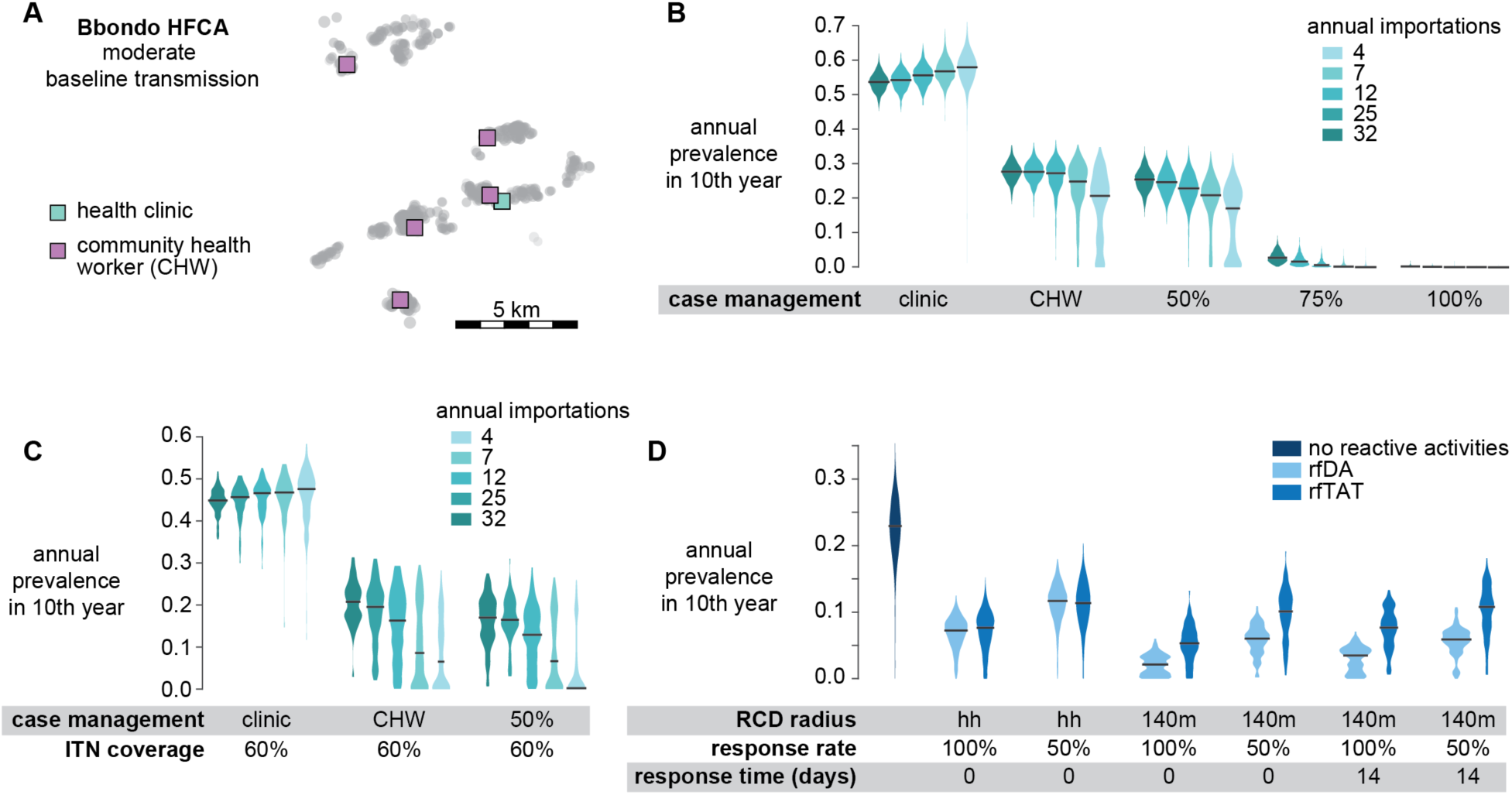
Preventing re-establishment of malaria in a previously moderate-transmission area requires good case management. (A) Households (circles) and locations of malaria treatment access points (squares) in Bbondo HFCA. (B) Prevalence in Bbondo 10 years post-elimination depends on case management rate but not importation rate. Case management rates for clinic-based and CHW-based case management are defined in Methods. No other interventions are simulated in Bbondo. (C) When case management rates are too low, maintaining regular ITN distributions lowers prevalence but does not prevent resurgence altogether. (D) Reactive case detection with focal drug administration (rfDA) or focal test and treat (rfTAT) can help maintain low prevalence when case management alone is insufficient. RCD radius indicates whether RCD is distributed only to the index case household (hh) or also to neighboring households within 140m. Response rate indicates the fraction of treated, symptomatic cases that trigger a reactive response. Response time is the number of days between treatment of the index case and administration of rfDA or rfTAT. Shown: 50% case management, 12 annual importations, and no vector control. In panels B-D, gray lines indicate medians of mean annual all-age prevalence of any infection across 100 stochastic realizations.

Previous work in a different model found that importation rate determines whether reestablishment happens but not the extent of reestablishment (17), which we also observe in our system (Figure 2B). Our simulations predict that in Bbondo HFCA, case management rate is the most important factor governing the extent of reestablishment, and a 75% case management rate is sufficient to prevent resurgence. Because transmission in Bbondo was never extremely high, population immunity is moderate, and many malaria infections acquired by Bbondo residents will be symptomatic and thus detectable by a good surveillance system. Treatment of these infections reduces the development of sexual stage parasites that transmit to mosquitoes, while lack of treatment leads to high-density sexual stage infections and high infectiousness (21, 22). Maintaining minimal local transmission through excellent case management perpetuates low population immunity, increasing the symptomaticity of new infections in the long term. Primary case management is thus highly effective for preventing secondary infections in Bbondo.

At low levels of case management, such as in the scenario where only the clinic is operational and treatment-seeking is distance-dependent, malaria transmission resurges in the 10 years post-elimination to levels above the previous baseline. Over a longer time horizon, population immunity will reestablish and prevalence will re-equilibrate to baseline. While malaria completely reestablishes in our model for every simulated importation rate under low case management, we find that higher importation rate results in sooner reestablishment. At the 10-year mark, simulations under the highest importation rate are further along in reestablishing population immunity than simulations under the lowest importation rate, and thus prevalence is lower for scenarios with more importation.

When case management is below 75%, maintaining regular distributions of insecticide-treated nets (ITNs) lowers prevalence but generally fails to contain prevalence to very low or near-zero levels (Figure 2C). At low importation rate and 50% case management, ongoing vector control can help prevent reestablishment with at least 50% probability, but in other cases, reestablishment remains likely.

Reactive case detection (RCD), where treated symptomatic cases trigger case investigation near the index case’s household, can lower prevalence when case management alone is insufficient (Figure 2D). We simulated RCD scenarios under the conditions of 50% case management and 12 imported infections per year, where prevalence is around 25% in the absence of RCD. We find that under certain conditions, RCD can reduce prevalence to under 5%. When RCD happens at the household level, where only the household of the index case is investigated but neighbors are not, case investigation with reactive focal-test-and-treat (rfTAT), where RDT-positive individuals are treated with an antimalarial, is just as successful as reactive focal drug administration (rfDA), where individuals are presumptively treated. RCD at the household level reduces prevalence from 25% to 8-12% depending on the fraction of index cases that receive follow-up. Increasing the RCD radius to 140m of the index case, which is current protocol in Zambia (23), can reduce prevalence to as low as 4%, as long as the response rate is high and CHWs can respond quickly. At 140m, rfDA is more successful than rfTAT, while at the household level, they are equally successful. In post-elimination Bbondo, the risk of asymptomatic infection is low (20), so the benefit of treating RDT-negative neighbors likely comes from the antimalarials’ short period of prophylactic protection.

### In areas with formerly high transmission, vector control and case management are both vital for limiting onward infection

Luumbo HFCA borders Lake Kariba (Figure 3A), and consequently transmission is much higher, with 42% of individuals testing positive for malaria by RDT during the dry season. Luumbo has around 12,000 residents living in roughly 744 households, but is geographically much larger than Bbondo and thus more sparsely populated. In addition to the clinic, 12 CHWs provide testing and treatment for symptomatic malaria, with one collocated with the clinic.

**Figure 3.**
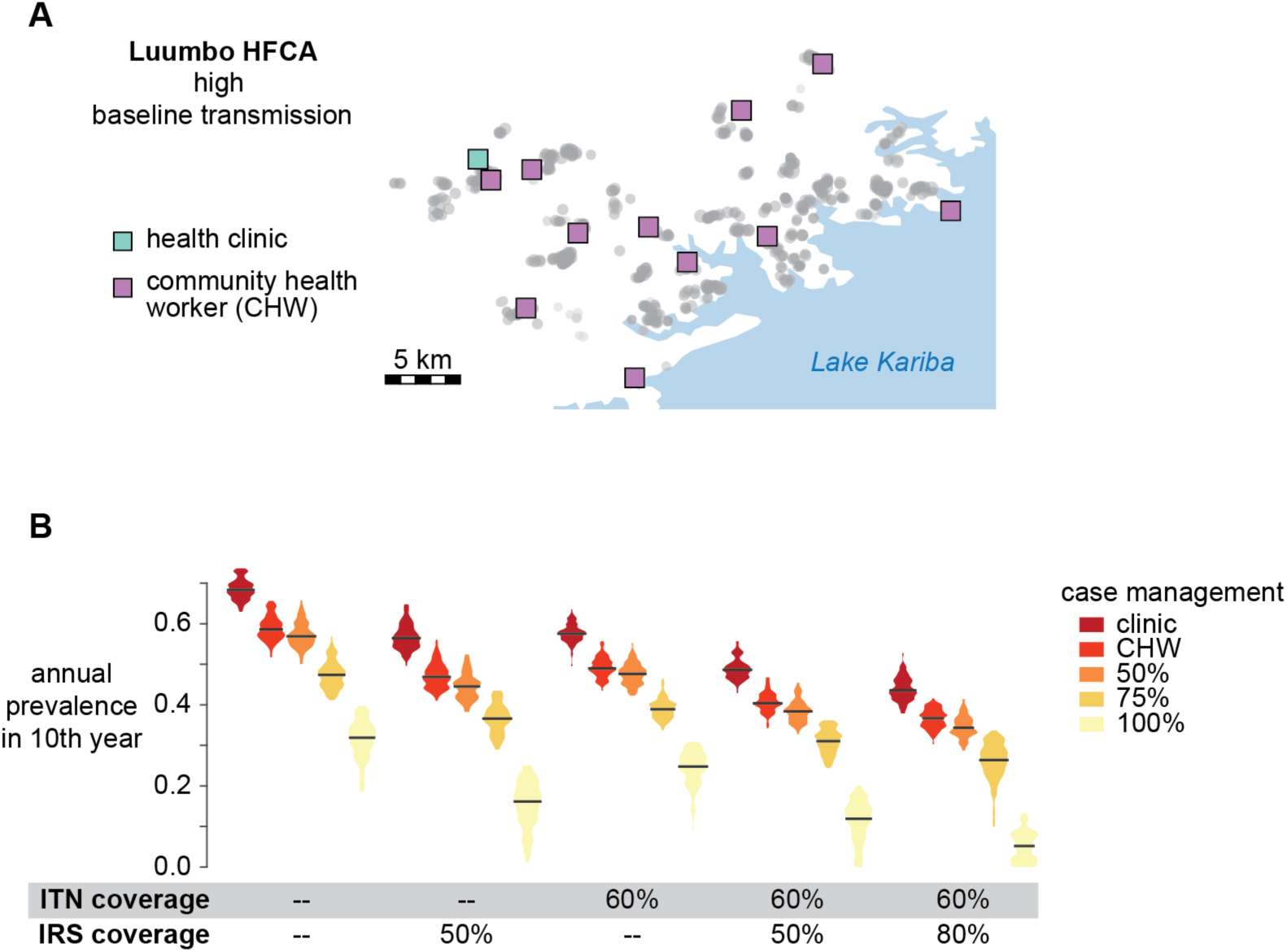
Maintaining excellent case management and routine vector control are vital for keeping prevalence low in areas with formerly high transmission. (A) Households (circles) and locations of malaria treatment access points (squares) in Luumbo HFCA. (B) Impact of case management and vector control on prevalence in Luumbo. No other interventions were simulated in these scenarios. Importation rate was on average 20 per year. Gray lines indicate medians of mean annual all-age prevalence of any infection across 100 stochastic realizations.

In post-elimination Luumbo, excellent case management and routine vector control can maintain low but not very low prevalence (Figure 3B). Unlike in Bbondo, 75% case management rate is insufficient to maintain near-zero prevalence in Luumbo, even with regular refreshment of ITNs and annual application of IRS. Scenarios that included both ITNs and IRS resulted in lower prevalence than scenarios with a single form of vector control.

### Reactive strategies can efficiently reduce prevalence in formerly high-transmission areas

We tested reactive strategies based on distribution of antimalarial drugs or IRS on top of routine ITN refreshment and 100% case management in Luumbo. As observed in Bbondo HFCA, rfDA and rfTAT can reduce prevalence from 12% to below 5% (Figure 4A). Unlike in Bbondo, rfDA is always superior to rfTAT in Luumbo, likely because of the substantial population immunity in Luumbo and preponderance of subpatent infections. Extending the radius of reactive focal activities to 140m is beneficial and achieving prompt response at high rates continues to be important for maximizing the impact of these activities.

**Figure 4.**
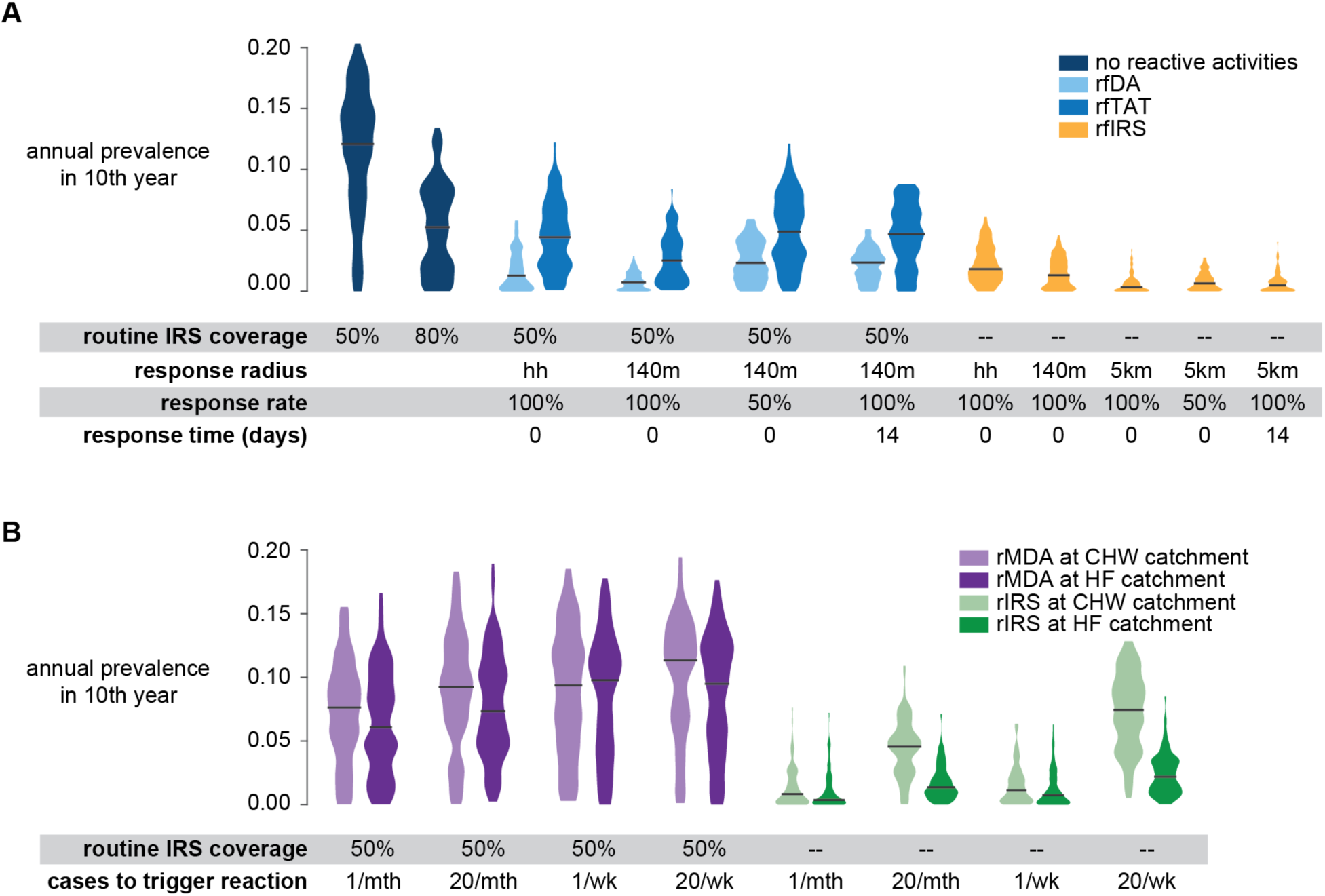
Ability of reactive interventions to maintain low prevalence in Luumbo HFCA. (A) Prevalence 10 years after elimination under various reactive focal intervention scenarios. (B) Prevalence 10 years after elimination under reactive intervention scenarios where cases are counted at the CHW or health facility (HF) level and reactive interventions are distributed to all households in the CHW or HF catchment respectively. Reactive interventions are delivered the day after the response threshold is reached. (A-B) All scenarios include 100% case management, 60% routine ITN coverage, and 20 importations per year. Gray lines indicate medians of mean annual all-age prevalence of any infection across 100 stochastic realizations.

Very low prevalence can be maintained in Luumbo when an index case triggers reactive focal IRS (rfIRS), where the index case’s household or neighborhood is sprayed with insecticide (Figure 4A). The rfIRS scenarios do not include annual HFCA-wide IRS spraying, and we limit each household’s eligibility for IRS to a maximum of once every six months. The impact of rfIRS increases with increasing radius of the focal activities, and rfIRS with a 5km radius around the index case household can keep prevalence below 1%. Reducing the response rate or increasing the response time still results in very low prevalence under this intervention scenario.

Rather than following up on each index case as it occurs, programs can also consider a reactive intervention scheme where index cases are tallied. If the total number of cases over a certain period exceeds a threshold, interventions are deployed. Using this framework, we consider reactive mass drug administration (rMDA), where all individuals are presumptively treated with antimalarials, and reactive IRS (rIRS) as possible response interventions (Figure 4B). We consider index case monitoring and intervention deployment at either the CHW-catchment scale or over all of Luumbo HFCA at once.

Simulations predict that rMDA is ineffective at maintaining low prevalence in Luumbo HFCA, likely because individuals missed during rMDA continue to transmit malaria. High coverage is difficult to achieve during MDA due to absent individuals, contraindications, and/or refusals (24, 25). In contrast, rIRS is predicted to have a much larger impact. When rIRS is responding to a single case observed over a week or month, monitoring and reacting at the CHW level is very nearly as successful as monitoring and reacting over the entire HFCA. If the response threshold is higher, at 20 symptomatic cases per week or month, monitoring and responding HFCA-wide is preferable.

IRS is an expensive intervention requiring trained personnel (26) and thus its use should be minimized when possible. WHO currently recommends focal IRS in elimination areas to target outbreaks (27). We counted for each routine IRS, rfIRS, and rIRS scenario how many houses were sprayed over the 10 years of simulation (Figure 5). Efficient IRS interventions are those that minimize both prevalence and houses sprayed. We find that rIRS is more efficient than rfIRS or routine IRS. Although rfIRS at a 5km radius resulted in the lowest prevalence, it also sprayed the most houses, on par with routine annual IRS at 80% coverage. rIRS can achieve nearly the same low prevalence while spraying over 10-fold fewer houses. While rIRS at the HFCA-level results in lower prevalence than rIRS at the CHW catchment level, monitoring and responding at the CHW catchment level can further halve the number of houses sprayed while continuing to maintain prevalence at or below 1%.

**Figure 5.**
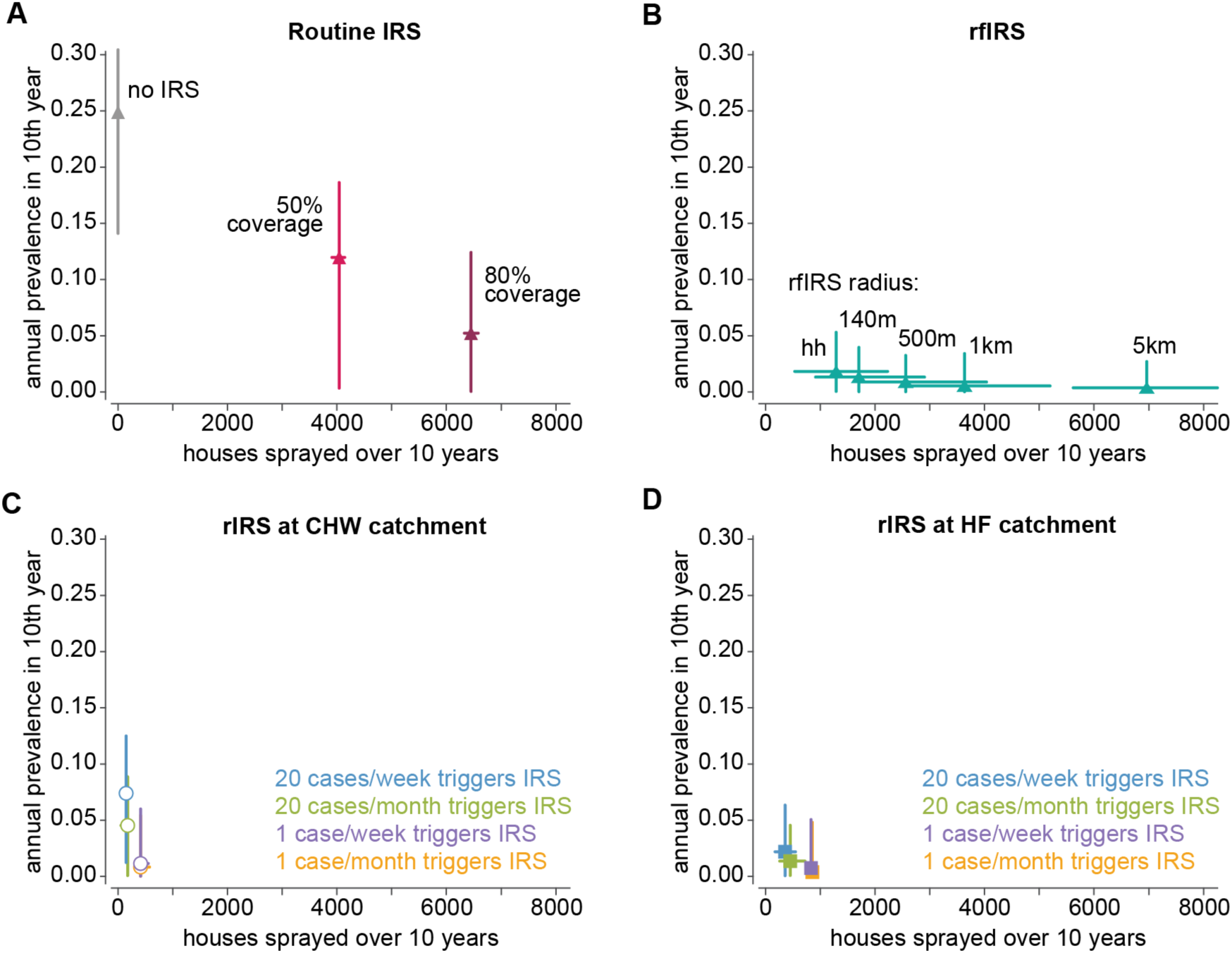
Number of houses sprayed under routine IRS, rfIRS, and rIRS and resulting prevalence in the tenth year after elimination. For rfIRS, no delay in response was simulated, and 100% of index cases received response. All scenarios include 100% case management, 60% routine ITN coverage, and 20 importations per year. Points indicate medians and lines indicate 95% observed range of mean annual all-age prevalence of any infection and number of houses sprayed across 100 stochastic realizations.

Several factors are key to the success of rIRS at the CHW level (Figure 6). Case management both reduces onward infection by clearing individuals of parasites and results in an index case recorded by surveillance systems. Without good case management, reactive strategies in Luumbo HFCA are badly hampered and insufficiently responding. Coverage of the rIRS operation also impacts the level of prevalence that is maintained, but not to the same degree as case management rate. The most effective rIRS strategy is to respond to a single symptomatic case anywhere in the HFCA by spraying the entire catchment, although responding at the CHW catchment to one case per week or one case per month is also highly effective. Unlike reactive focal strategies, the delay to respond matters less in catchment-based reactive strategies, because the scale of response is much larger and infections that have spread beyond the household or 140m radius can still be contained.

**Figure 6.**
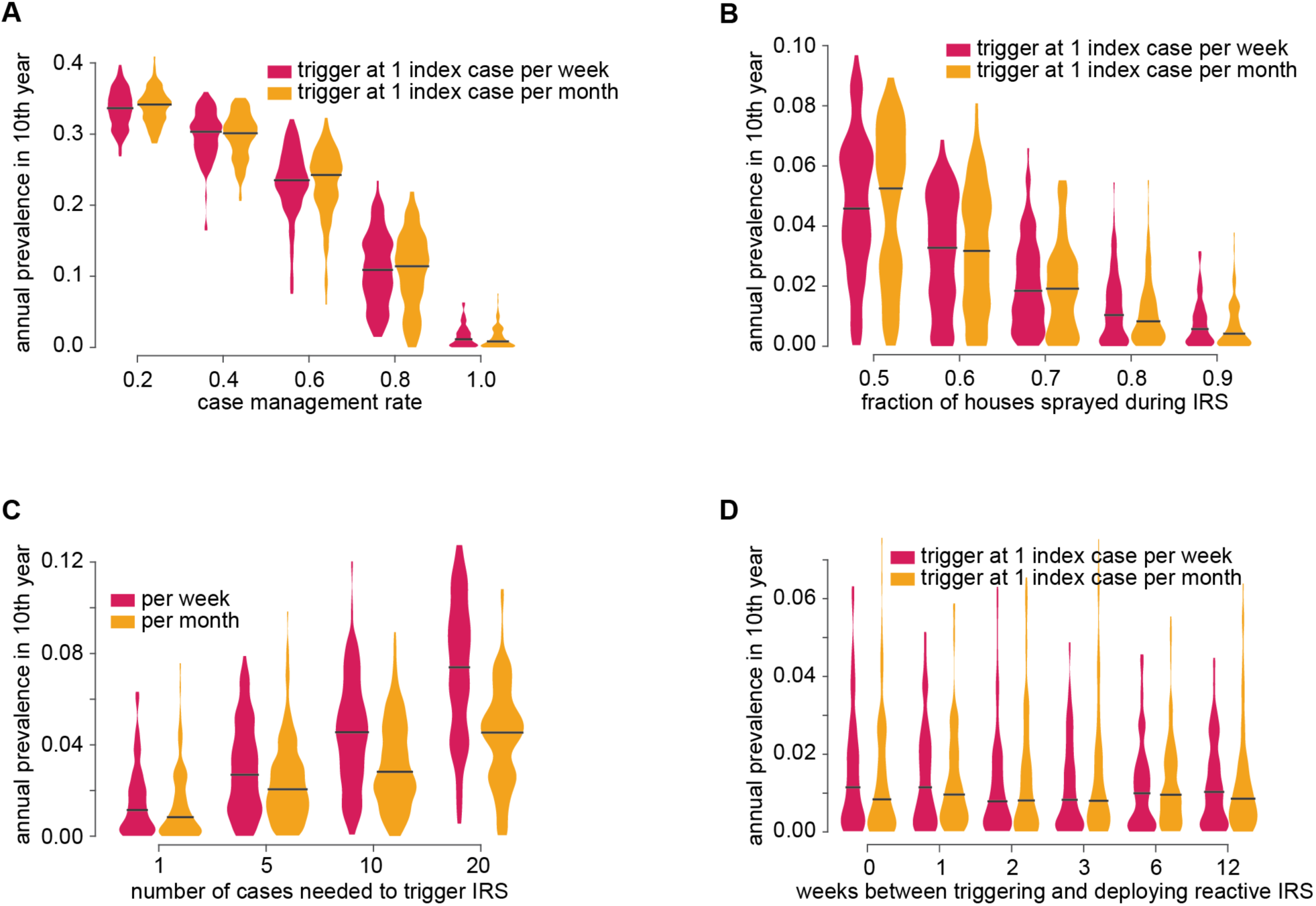
Dependence of reactive IRS impact on (A) case management rate, (B) IRS coverage, (C) number of cases needed to trigger IRS, and (D) response time. All scenarios include 100% case management, 60% routine ITN coverage, and 20 importations per year. All scenarios are monitoring and reacting at the CHW catchment level. Default scenario parameters used unless otherwise indicated: case management 100%, rIRS coverage 90%, 1 case triggers rIRS, no delay in rIRS response. Gray lines indicate medians of mean annual all-age prevalence of any infection across 100 stochastic realizations.

## Discussion

In areas of formerly moderate and formerly high transmission, good case management is the foundational intervention to contain the spread of imported infections. In our model framework, other interventions could not completely compensate for inadequate case management. This finding is well-aligned with previous work in different models (17). As models have also predicted that good to excellent case management is required for reaching elimination in the first place, it is critical that countries build their health systems sustainably such that treatment and reporting rates for symptomatic malaria remain universally high for many years.

Treatment of symptomatic cases both prevents onward transmission and provides critical information for routine surveillance systems, particularly in this context for outbreak response. For countries where individuals often seek care in the private sector, surveillance becomes an even more complex challenge. If case reporting is not integrated across all sectors, response rates of reactive interventions may become too low to contain transmission. Other models have also identified the response rate of reactive activities as a critical factor for success (17, 28).

As time since elimination increases, excellent case management increases the impact of case management in later years. When most symptomatic cases are treated, individuals have less opportunity to develop immunity to malaria symptoms, and the population as a whole becomes less immune to symptoms. In addition to population cycling bringing new malaria-susceptible infants to the population and removing semi-immune adults, immunity to malaria may wane within each individual (29, 30). However, this waning is little-understood, particularly how waning immunity to symptoms may differ from waning immunity to high-density infection, which could lead to more infectious individuals. In this study, we have assumed no waning of either type of immunity. If waning immunity to symptoms is actually fast and substantial, then we have underestimated the impact of case management; conversely, if waning immunity to high-density infections is fast and substantial, then preventing reestablishment may be more challenging than predicted.

RCD with antimalarial drugs has previously been shown to have only a minimal or moderate impact on reaching elimination (31-34). Post-elimination, we find that RCD can help sustain a near-eliminated state, but only under certain performance criteria. The key difference between RCD to reach elimination and RCD to prevent resurgence is the extent to which asymptomatic infections exist outside the radius around the index case. On the road to elimination, asymptomatic infections may be common even in areas outside the RCD radius of the index case, simply because transmission was happening across the area. Since RCD is not deployed if there is no index case nearby, the impact of RCD is limited in this context. After elimination, infections are more likely to be clustered around a single point, the point of reintroduction, and a larger fraction of asymptomatic infections would thus be targetable by RCD.

While routine interventions such as case management and maintaining ITN coverage are necessary, our simulations also suggest that post-elimination, some interventions can instead be deployed reactively. In our framework, reactive IRS is both more effective and sprays fewer households, which could lead to cost savings. Operationally, reactive IRS could be more challenging to implement than routine IRS because where and how much will be needed is difficult to predict. However, we also find that even when there are substantial delays in IRS response, this intervention is still able to contain transmission. Indeed, reactive IRS may also make sense as an intervention on the path to elimination as well, since even within an HFCA there can be substantial variation in transmission intensity.

Our simulations predict that reactive IRS can successfully maintain very low prevalence at multiple scales of reaction, although the number of houses sprayed differs by over an order of magnitude. An important source of uncertainty in this analysis is the flight patterns of local vectors, which will influence the appropriate spatial scale of any outbreak response activity. Literature suggests that anopheles vectors fly as much as necessary between blood meals and breeding sites (35), and this distance can vary substantially between settings (36). We have assumed vectors mix well within 500m, and less so up to 3km. Under these assumptions, responding at the CHW catchment is sufficient to contain transmission while limiting number of houses sprayed. In Luumbo, a CHW is responsible on average for an area of over 38km^2^, although catchments vary widely in size. If additional CHWs are recruited, the CHW catchment may no longer be sufficiently large for adequate outbreak response to be limited to this area.

In this work, we have assumed that baseline transmission potential, modeled as vector larval habitat availability, does not change from year to year. This may be a pessimistic assumption as economic development, leading to housing improvement and environmental management, has historically driven down malaria transmission (37). Climate change could also affect local transmission potential in the long term, although predicted increase in malaria risk has large uncertainties (38, 39). It is unclear which long-term trend will dominate in Bbondo and Luumbo HFCAs.

Maintaining near-zero infection in areas with high potential for malaria transmission, in the face of continual importation of new cases, will be a challenge. Malaria surveillance and operations may need to continue for years even though local burden is minimal. Sustainable, high-quality interventions are key to success, particularly excellent access to care for symptomatic malaria. Building on this foundation, programs can consider how to redesign routine interventions such as IRS into rapid and agile outbreak response operations. Other routine vector control may need to continue indefinitely until importations cease, particularly if transmission was high prior to elimination efforts. Above all, comprehensive and integrated surveillance is essential to identify outbreaks quickly and trigger response interventions.

## Methods

### Simulation framework

A previously-developed spatial model of malaria transmission in southern Zambia (20) was adapted for use in this study (Figure 1A). This model is an agent-based patch model(40) where each patch represents a single household, and both humans and mosquitoes can move between households. Areas between households were not modeled. Vector life cycle and feeding behavior, within-host parasite dynamics, and acquisition of immunity were parameterized according to field observations and described elsewhere (41-43). Dominant vector species were *Anopheles arabiensis* and *Anopheles funestus*, with *arabiensis* predominating, and 50% and 95% indoor biting respectively.

We selected two of the areas described in (20), Bbondo and Luumbo health facility catchment areas (HFCAs), and modified each as follows. We froze the simulation state for each HFCA on January 1, 2017, including each human’s immune status and intervention history. We next artificially removed all parasites from every infected human and mosquito, creating a pair of HFCAs where malaria has been instantaneously eliminated. We then carried forward each simulation for 10 years, allowing residents to travel to a generic uneliminated area with a forced, seasonally-varying entomological inoculation rate (EIR) (Figure 1B). Residents returned to their home HFCA possibly infected with malaria. Mosquitoes could not travel between the HFCA and the uneliminated area. During the 10 simulation years, we considered various intervention packages in the home HFCA, and in the 10^th^ year we measured the all-age true prevalence of any malaria infection, averaged over the year (Figure 1C). Each intervention scenario was run for 100 stochastic realizations.

Vectors were modeled as individual agents with distance-dependent movement between households, with 90% of vector flights within 370m, 97% within 500m, and small probability of movement to households up to 3km distant (20). Humans could move within their home HFCA for overnight stays lasting on average 3 days; individuals made on average 0.075 overnight trips per person per year. Human trips to the uneliminated area lasted on average 30 days to reflect local patterns of movement to lakeside areas for farming and fishing. Bbondo and Luumbo residents had the same probability of making a trip to the uneliminated area, although because Luumbo’s population was 12,000 compared with Bbondo’s 8,000, more total trips were made in the Luumbo model and the number of imported infections each year was higher. The force of infection in the uneliminated area was sampled evenly in log space from 2 infectious bites per person per year to 54 infectious bites per person per year. Unless otherwise specified, scenarios were run with 8 infectious bites per person per year in the uneliminated area, resulting in on average 12 imported infections per year in Bbondo and 20 in Luumbo (S1 Figure).

### Interventions

We considered the following interventions in Bbondo and Luumbo HFCAs.

Case management for symptomatic malaria: individuals experiencing clinical malaria received artemether-lumefantrine (AL). Pharmacokinetics and pharmacodynamics of AL were calibrated to clinical study data (44). At 50%, 75%, and 100% case management rates, all individuals sought care with the specified probability for both uncomplicated and severe malaria. At “clinic” and “CHW” levels of case management, individuals sought care according to their household’s distance from the clinic or the nearest community health worker (CHW), whose locations are shown in Figure 2A and Figure 3A for Bbondo and Luumbo respectively. The distance- and age-dependence of care-seeking was informed from surveys conducted in this area (11, 19, 20, 45). Children under age 15 sought care at a relative 50% higher rate than did adults, and individuals with severe malaria were 80% relatively more likely to seek care than individuals with uncomplicated malaria. These factors combine to create overall case management rates of, in Bbondo, 22% with the clinic alone and 44% with CHWs; in Luumbo, 17% with the clinic alone and 39% with CHWs.

Routine distribution of insecticide-treated nets (ITNs): mass distribution of ITNs to individuals of all ages every 3 years in the middle of the dry season. ITNs were distributed at 60% coverage at the individual level, and individuals who owned a net were assumed to use it every night. Coverage was not correlated across distribution events or within households. ITN efficacy was modeled identically for both vector species, with initial killing and initial blocking rates of 0.6 and 0.9 respectively, each decaying exponentially with half-lives of 1.4 and 2.7 years respectively. To replicate observed behaviors in this region, individuals discarded ITNs with mean retention time of 9 months.

Routine administration of indoor residual spraying (IRS): annual IRS campaigns conducted immediately prior to the wet season. Probability of a household receiving IRS was not correlated between campaigns. IRS efficacy was identical for both vector species, with initial killing rate of 0.6 and half-life of 9 months.

Reactive case detection (RCD) with focal response: A treated symptomatic index case triggered case investigation, which could take the form of reactive focal test-and-treat (rfTAT) with a rapid diagnostic test (RDT), where individuals who test positive receive AL; reactive focal drug administration (rfDA), where all individuals receive AL; or reactive focal IRS (rfIRS), where households or neighborhoods receive IRS. For rfTAT and rfDA, 60% of individuals were assumed to be at home and able to take antimalarials. No limit was set on minimum duration between rfTAT or rfDA events in a single household. We considered rfTAT and rfDA at either only the index case household or the index case household and all neighboring households within 140m. For rfIRS, 90% of households were assumed to accept and receive IRS, and a household could be sprayed at most once every 6 months. rfIRS was efficacy parameterized as above for routine IRS. During rfIRS, no rfTAT or rfDA was simulated, and we considered rfIRS at only the index case household or the index household and neighbors within 140m, 500m, 1km, and 5km.

Monitoring and reactive response at the catchment level: Number of treated cases was tallied at either the CHW catchment or HFCA level. CHW catchments were defined by assigning each household to its closest CHW. Upon reaching a threshold number of cases over a given duration, a response was triggered over the area of surveillance. We considered reactive IRS (rIRS) or reactive mass drug administration (rMDA) responses. rIRS was parameterized as above for rfIRS. During rMDA, all covered individuals received DHA-piperaquine (DP), two rounds of distribution were conducted separated by 1 month with independent coverage between rounds, and coverage was modeled at 70%. Areas that received rIRS or rMDA were ineligible for another distribution of rIRS or rMDA for a period of 6 months. We considered the threshold number of cases, the duration over which to count cases, the geographical area of monitoring and response, the response coverage, the delay period between reaching the threshold and responding, and the underlying case management rate as factors impacting the success of these interventions.

## Acknowledgements

The authors thank Benoit Raybaud, Bryan Ressler, and Svetlana Titova for software support, and the Malaria Modeling Consortium and Kim Lindblade for helpful feedback.

## Supporting information

**S1 Figure.**
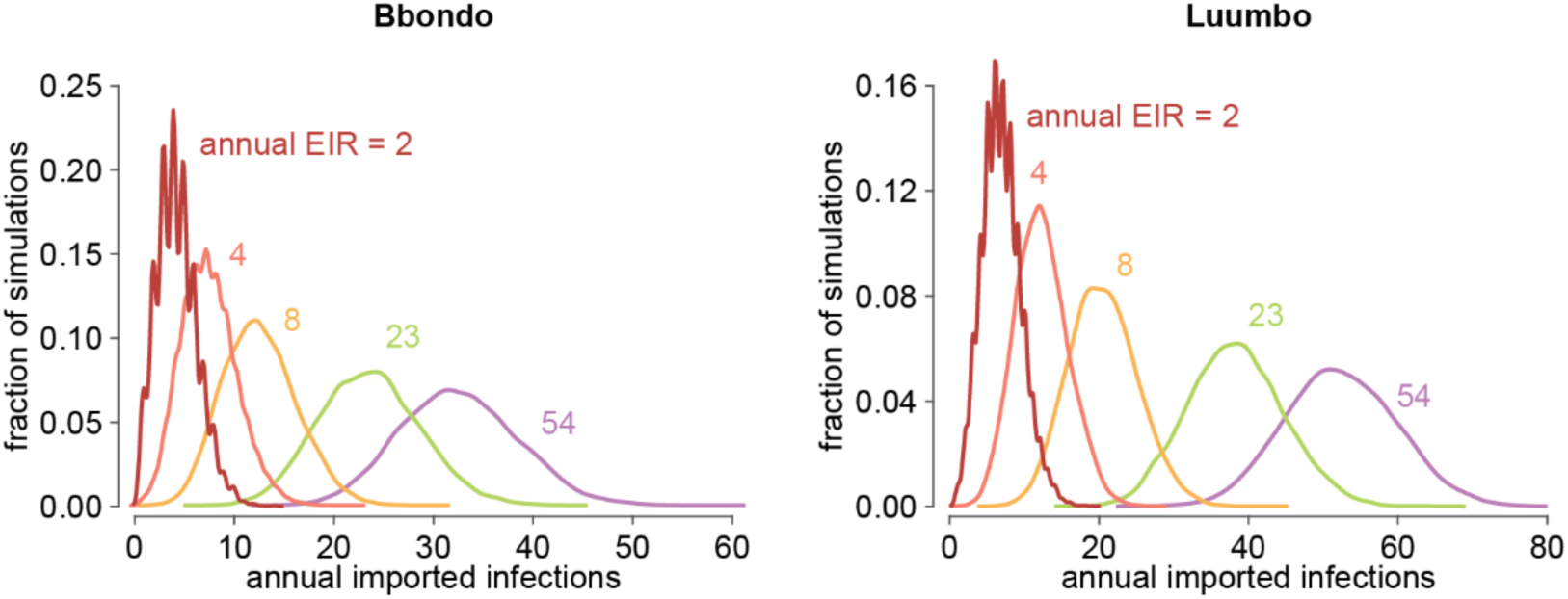
Distributions of number of imported infections per year, in each of Bbondo and Luumbo HFCAs, at the five levels of transmission intensity (entomological inoculation rate (EIR) measured in infectious bites per person per year) considered in the uneliminated area to which Bbondo and Luumbo residents may travel.

